# Geographical Distribution of *Biomphalaria* snails in East Africa

**DOI:** 10.1101/2021.11.04.467236

**Authors:** Victor O. Magero, Sammy Kisara, Christopher M. Wade

## Abstract

There is limited information on the distribution of *Biomphalaria* snails, an important snail intermediate host of schistosomiasis, in East Africa. This study assessed the incidence and geographical distribution of *Biomphalaria* snails in Kenya, Uganda and Tanzania. Maximum Entropy modeling was used to predict the potential distribution of *Biomphalaria* snails, in line with habitat suitability. Malacological surveys were then conducted guided by MaxEnt predictions and information obtained from previous research papers. The surveys were conducted at a total of 156 sites including streams, rivers, lake shores, dams and irrigation canals over a 3-year period (2018 to 2020). Geographical coordinates, ecological and physicochemical information was recorded for the sites visited. Snails were identified morphologically, based on shell characters using established identification keys. *Biomphalaria* snails were found at 23.07% (36/156) of the sites sampled. Streams proved to be the habitats most preferred by *Biomphalaria* snails (50% of all of the sites where the snails were found were streams), followed by rivers (20.6%), irrigation canals (8.8%), lake shores (8.8%), springs (5.9%), and dams (5.9%) with snail abundance increasing with increase in temperature and decrease in water depth. *Biomphalaria* snails were found in the Lake Victoria basin, Mwea Irrigation Scheme and Eastern Province of Kenya, the Lake Albert region, Lango region, Soroti district, Lower Moshi irrigation scheme, Babati district, Iringa region, Tabora region and Kigoma region. Information on the distribution of *Biomphalaria* snails in East Africa will aid in developing prevention and control strategies for schistosomiasis.

**AUTHOR SUMMARY:** Few studies have been conducted on the distribution of *Biomphalaria* snails in East Africa with previous studies mostly isolated projects restricted to single divisions, districts and regions. Knowledge on the distribution of snail intermediate hosts can be helpful in establishing schistosomiasis transmission surveillance systems for detecting emerging and prevailing incidences of schistosomiasis. We undertook malacological surveys of freshwater sites across Kenya, Uganda and Tanzania. A total of 156 sites were sampled and *Biomphalaria* snails were found at 36 of the sites. Streams yielded the highest number of snails, in comparison to the other habitats that were sampled. Temperature and water depth were established to be statistically significant ecological and physicochemical factors that influence incidences and abundance of the snails. This study provides important information on the distribution of an important snail intermediate host in East Africa and the knowledge obtained herein can be helpful in establishing appropriate schistosomiasis control initiatives.

## INTRODUCTION

Schistosomiasis is a neglected tropical disease caused by flatworms of the genus *Schistosoma*. The disease is ranked second to malaria in terms of tropical diseases of public health importance (Colley *et al.*, 2014). It is estimated that about 1 billion people globally are at risk of being infected with the disease and about 250 million people are currently infected (Gryseels, 2012). 200,000 deaths are attributable to the disease annually (Colley *et al.*, 2014). The disease has debilitating effects on individuals because it renders them unproductive due to malaise and disability. In particular, the disease is catastrophic to school aged children because it impedes them from attending school due to physical incapacity associated with the disease, thus jeopardising their chances of having a brighter future. There are two forms of schistosomiasis, urogenital schistosomiasis caused by *S. haematobium* and intestinal schistosomiasis caused by *S. mansoni* and *S. japonicum* (Colley *et al*., 2014). Other human *Schistosoma* parasites albeit not significant from a public health perspective include *S. intercalatum, S. mekongi* and *S. guineensis* (Gryseels, 2012). *Schistosoma mansoni* is responsible for most cases of human intestinal schistosomiasis in Africa and the Americas; it is solely responsible for 83 million out of an estimated total 250 million global schistosome infections (Morgan *et al.*, 2005).

Snail intermediate hosts are important because they support the transformation of *Schistosoma* parasites into infective stages (Zeng *et al*., 2017). Snail intermediate hosts for *S. mansoni* belong to the genus *Biomphalaria*. In the Americas, intermediate hosts *for S. mansoni* include *B. glabrata, B. tenagophila* and *B. straminea* (Habib *et al*., 2021). Fossil records assert that African *Biomphalaria* species originated from *B. glabrata*, during the Plio-Pleistocene period when *B. glabrata* invaded the African continent (Campbell *et al*., 2000). In East Africa, there are several species of the genus *Biomphalaria* including *B. sudanica*, *B. choanomphala, B. smithi, B. stanleyi, B. angulosa* and *B. pfeifferi* (Standley & Vounatsou, 2012).

*B. pfeifferi, B. sudanica* and *B. stanleyi* have been identified from Lake Albert (Rowel *et al*., 2015) while *B. sudanica* and *B. choanomphala* have been identified from Lake Victoria (Standley *et al*., 2011; Standley *et al*., 2014). *B. sudanica* and *B. smithi* have been found from Lake Edward (Mandahl, 1957). *Biomphalaria* snails, although not identified to the species level have been collected from Mount Elgon, Rwenzori Mountains and Fort Portal crater lakes (Stanton *et al*., 2017). *B. sudanica* snails have also been found in Northern and North Eastern parts of Uganda (Habib *et al*., 2021). *B. pfeifferi* snails have been found in Uganda at Lwampanga, Masindi Port, Muzizi, Ngamilajojo, Butiaba and Ntoroko (Lake Albert) (Jorgensen *et al*., 2007). *B. pfeifferi* snails have also been found in Northern and North Eastern parts of Uganda (Habib *et al*., 2021). In Kenya, *B. pfeifferi* snails have been found at Muthamo seepage, Matingani seepage, Mbondoni dam, Kangonde dam, Mwea East, Mwea West, Onsando dam, Grogan Canal, Kamayoga stream, Asawo stream, Kwahoma stream and Martin’s drain (Bandoni *et al*., 1990). Malacological surveys by Opisa *et al*. (2011), at informal settlings of Kisumu City, Kenya identified two *Biomphalaria* species, *B. sudanica* and *B. pfeifferi*, revealing a risk of schistosomiasis transmission at this location. *B. pfeifferi* snails were also found in Kenya, at the Mwea irrigation scheme, Asawo stream and Mukou stream by Mutuku *et al*. (2020). A study by Buddenborg *et al*. (2017) identified and collected *B. pfeifferi* from Kasabong stream, Asembo village (Western Kenya). *B. pfeifferi* snails have also been reported from Kenya at Kibwezi (Jorgensen *et al*., 2007) as well as Chebunyo dam (Bomet County) and Churo pond (Kirinyaga County) (Bandoni *et al*., 2000). *B. pfeifferi* snails have also been collected from Asawo River and Kasabong stream, in Western Kenya (Lu *et al*., 2016). In Tanzania, *B. pfeifferi* snails have been found at the southern part of the country, around Gombe National Park (Bakuza *et al*., 2017) and have been identified and collected in the Northern part of Tanzania (in Babati district and Lower Moshi irrigation scheme) by Lydig (2009) and Kisanga (1991) respectively. *B. pfeifferi* has also been found in Kilombero district, Tanzania (Utzinger & Tanner, 2000). The previous studies on the distribution of *Biomphalaria* snails in East Africa (Kenya, Uganda and Tanzania) have not covered expansive geographical areas. The greater the abundance of snail intermediate hosts in any geographical location, the higher the probability that human beings will come in contact with *Schistosoma* parasites and be infected (Bakuza *et al*., 2017).

Intestinal schistosomiasis has been reported to be endemic in several localities in East Africa (Aula *et al*., 2021). However, there has been little information on the role that *Biomphalaria* snails play in transmission of schistosomiasis in those localities. An increase in schistosome infections in snail intermediate hosts, leads to contamination of water bodies with active cercariae thus meaning that human beings become at increased risk of being infected with the parasites. Some of the localities and regions where intestinal schistosomiasis has been reported to be endemic, with little to no reports on snail intermediate hosts include Lango region (Adriko *et al*., 2018), West Nile region (Imran, 2014), Soroti district and Kibimba irrigation scheme (Ejotre *et al*., 2014). There is a therefore a need to extensively establish the distribution of *Biomphalaria* snails in East Africa, because of limited information about their distribution and their significance as an important intermediate host in the transmission of schistosomiasis. Understanding the distribution of *Biomphalaria* snails in East Africa would influence decision making as far as development of effective and integrated strategies to combat transmission of human intestinal schistosomiasis is concerned.

This study therefore aimed to assess the incidence of *Biomphalaria* snails and their current geographical distribution in East Africa as well as to determine physicochemical and ecological parameters that are associated with the abundance of the snails.

## MATERIALS AND METHODS

### Ecological modeling and evaluation

Ecological modeling was done using MaxEnt (Phillips *et al*., 2006), a machine learning algorithm that is used to predict the potential distribution of species, based on the principle of maximum entropy in a situation where there is incomplete information about the distribution of a given species and the environmental factors that promote its habitation. To conduct maximum entropy species distribution modelling, occurrence points-data (sites where *Biomphalaria* have been found previously) and environmental layers (bioclimatic factors and environmental factors that affect the distribution of species) are required.

Occurrence records for *Biomphalaria* snails were obtained from the Global Biodiversity Information Facility database (GBIF (https://www.gbif.org/). A total of 56 occurrence records were retrieved from GBIF. Bioclimatic variables were obtained from WorldClim.org and soil type and land cover variables obtained from respective databases. Soil type variables were derived from https://www.isric.org/explore/soil-geographic-databases and land cover variables were derived from http://2016africalandcover20m.esrin.esa.int/download.php

### Malacological surveys

MaxEnt predictions, past knowledge on the distribution of *Biomphalaria* snails and areas where intestinal schistosomiasis has been reported to be endemic in East Africa guided where we undertook our malacological surveys. Additionally, we visited water bodies in regions that have previously been reported as having cases of intestinal schistosomiasis, on the basis that there must be snail intermediate hosts that transmit the disease in those localities.

Malacological surveys were conducted from January 2018 to March 2018, December 2018 to February 2019 and January 2020 to March 2020, from 156 sites, across East Africa (Figure 1). Sampling was carried out in the months of December, January, February and March because generally these are the months in East Africa where there are no heavy rains (overflowing water). Overflowing water is associated with high water currents which normally destroy habitats of *Biomphalaria* snails and sweep the snails downstream. Surveys included 75 sites in Kenya, 26 sites in Uganda and 55 sites in Tanzania. The surveys involved searching edges of streams, river banks, irrigation canals, lake shores, springs and dams for snails, for around 30 minutes, usually covering a distance of approximately 20m and an area of around 7m^2^ per site. Snails that were found were collected using a handheld scoop. Geographical coordinates for each site were recorded using a handheld Garmin eTrex 10 GPS (Kansas City, USA). The snails were then sorted by turning them upside down on the wire mesh of the collection scoop to check their shell characters. Identification keys developed by (Brown 1994; Kristensen,1987) were used to identify *Biomphalaria* snails based on their morphological characters including the general shape of the shell, the shape of the whorls, the number of coils and the shape of the aperture. The snails were transferred to a collection jar then washed and placed in Falcon tubes while in the field. To preserve the snails, absolute ethanol was added to the lab bottles containing the snails to a volume double that occupied by the snails. Ethanol in the lab bottles was changed two to three times, to ensure that ethanol penetrated into the snails’ tissues. After completion of fieldwork, the lab bottles containing the snails were stored in a - 20°C freezer.

**Figure 1:**
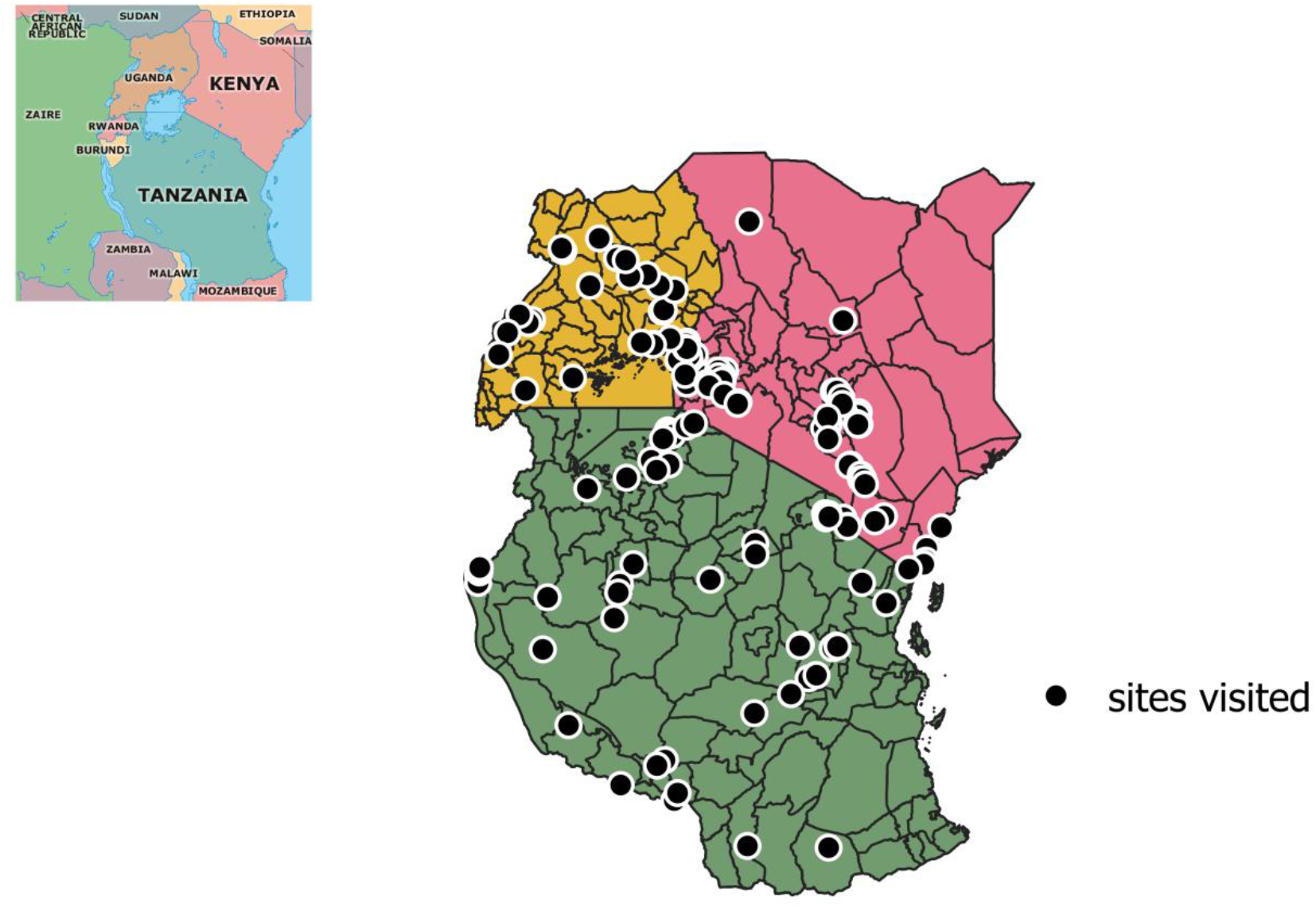
A map of East Africa showing sites that were visited for purposes of malacological surveys

### Ecological and physicochemical parameters

Ecological and physicochemical parameters associated with snail abundance were recorded. These parameters included vegetation type, water temperature and Ph. Water temperature and Ph were recorded using a portable water metre (Hanna Instruments, Møllevænget, Sweden).

Water depth was also recorded using a 1 metre ruler. Furthermore, water velocity was recorded using a flowmeter. Additionally, we noted and recorded the soil type for each site.

### Data processing and analysis

Maps showing sites that were visited, sites where *Biomphalaria* snails were found in relation to snail abundance were created by QGIS version 3.12.3. The name of the site, coordinates, availability of snails, snail numbers, ecological and physicochemical parameters for each site were recorded in Microsoft Excel.

For the purpose of determining ecological and physicochemical factors associated with snail abundance, the Gaussian log function was used. In this function, *Biomphalaria* snail abundance was used as an outcome while independent variables were the physicochemical and ecological parameters. Negative Binomial Generalised linear mixed models (GLMMs) were fitted in R Version 4.1.0 using the package ‘glmmTMB’ to test association between snail abundance and parameters.

## RESULTS

### 4.1 Potential distribution of *Biomphalaria* snails determined by MaxEnt

*Biomphalaria* snails are predicted to be widely distributed in East Africa based on MaxEnt analysis (Figure 2). Locations where there is a high probability that *Biomphalaria* snails will be found are shown in red while the blue colour shows locations where there is little probability that the habitat is suitable for the habitation of *Biomphalaria* snails. The figure suggests that *Biomphalaria* is endemic in East Africa.

**Figure 2:**
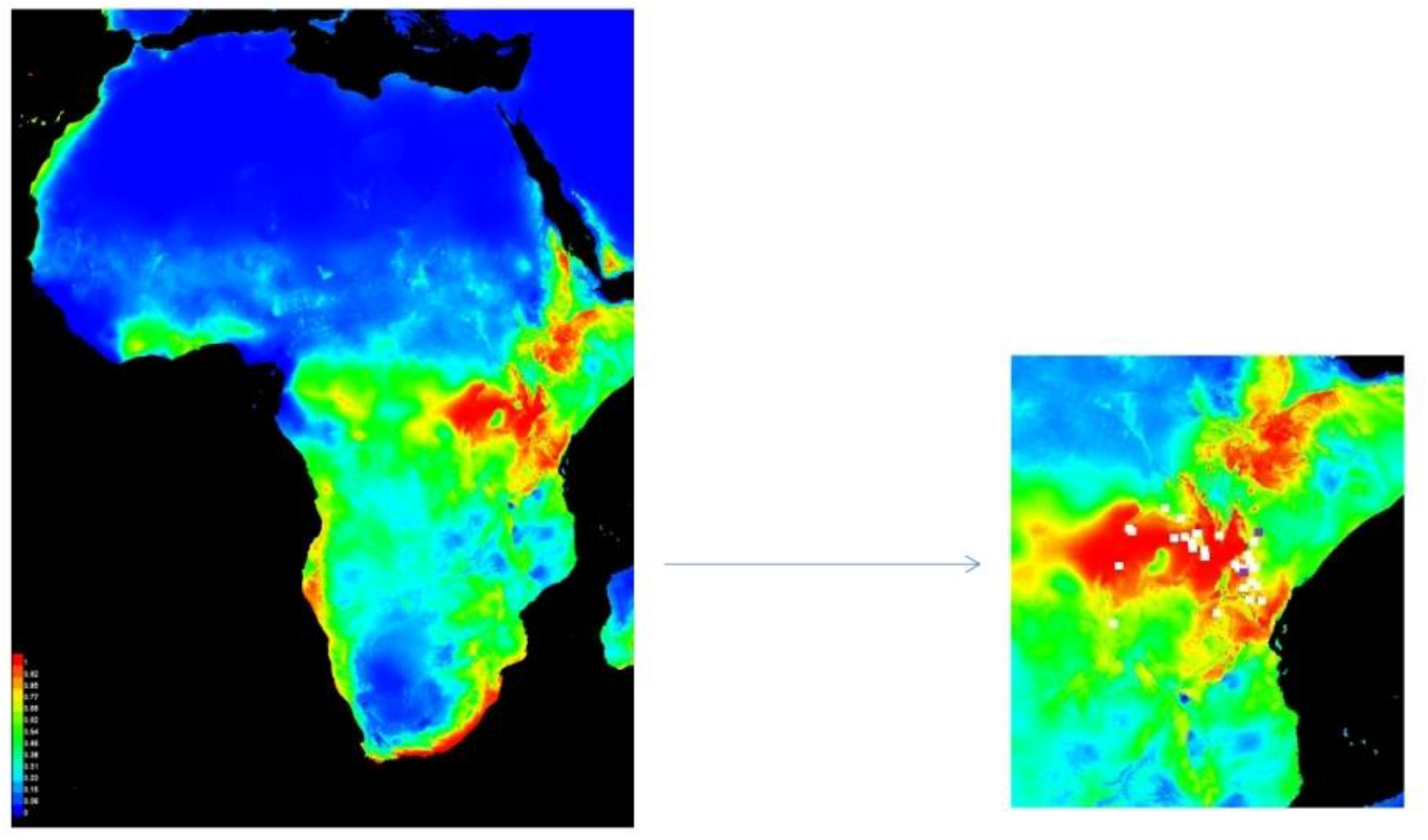
Potential distribution of *Biomphalaria* snails in Africa and East Africa, courtesy of MaxEnt

### 4.2 Presence and incidence of *Biomphalaria* snails

A total of 156 sites were visited across East Africa and *Biomphalaria* snails were found at 36 of these sites (Figure 3). *Melanoides tuberculata* was found at 7 of the 156 sites visited and even though the habitat of the sites where *M. tuberculata* were found were suitable for the habitation of *Biomphalaria* snails, *Biomphalaria* snails were only found together with *M. tuberculata* at one site (Ntorokom, Eastern Uganda).

**Figure 3:**
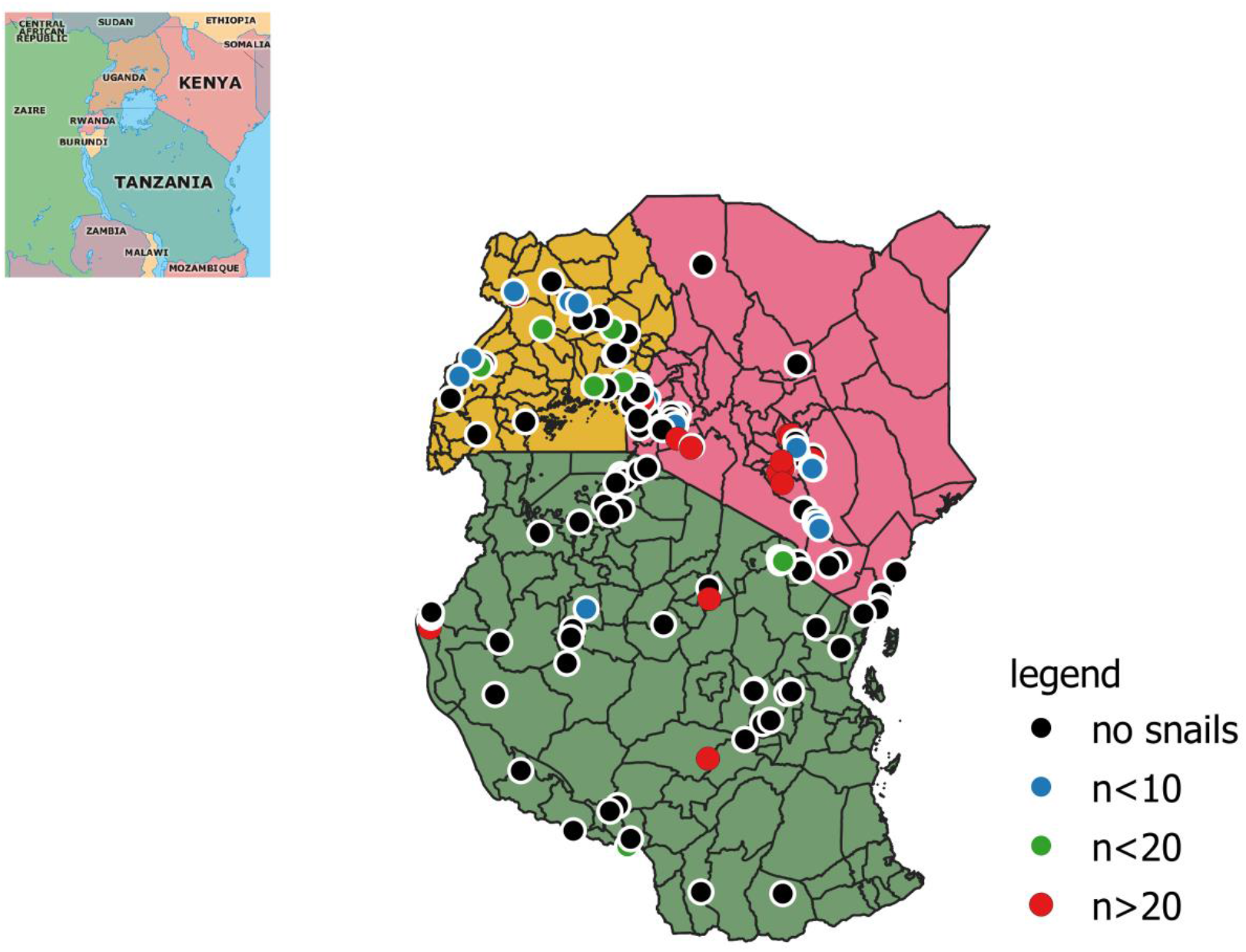
Incidences of *Biomphalaria* snails in East Africa, in regard to their abundance

*Biomphalaria* snails were found in rivers, streams, lakes shores, irrigation canals, springs and dams. Most of the sites where *Biomphalaria* were found were streams (50%), followed by rivers (20.6%), irrigation canals (8.8%), lake shores (8.8%), springs (5.9%), and dams (5.9%).

Supplementary Table 1 shows all sites that were visited for the purposes of conducting malacological surveys.

### 4.3 Ecological and physicochemical parameters

Data on vegetation availability, type of vegetation, water temperature, soil type, Ph, Water depth and water velocity was recorded for each site. Water temperature at the sites ranged from 19.1°C - 27.4°C and the median (interquartile range) water temperature was 23.6°C (21.18°C-25.23°C). The water-depth of the sites ranged from 19.3cm - 41.3cm and the median (interquartile range) water depth was 27.45 cm (23.1cm −32.4cm). Based on negative binomial regression in glmmTMB, only temperature and water depth had a statistically significant positive relationship with the abundance *Biomphalaria* snails (p-values of 0.00184 and 0.0187 respectively). The other variables (vegetation, algae, pH, soil type and presence of competitor snails) did not show any statistically significant relationship with the abundance of *Biomphalaria* snails (Table 1).

**Table 1:** The correlation between snail abundance ecological and physicochemical parameters courtesy of glmmTMB in R

| Variables | Estimate | Std. Error | Confidence Interval | p-value |
| --- | --- | --- | --- | --- |
| Intercept | -7.11562 | 3.93026 | -14.81878984 - 0.5875451 | 0.07022 |
| Vegetation | 1.82671 | 1.48377 | -1.08143734 - 4.7348507 | 0.21828 |
| Algae | 0.18526 | 1.10139 | -1.97341919 - 2.3439365 | 0.86642 |
| Temp | 0.2508 | 0.08051 | 0.09300019 - 0.4085942 | 0.00184 ** |
| Ph | 0.33459 | 0.43332 | -0.51471196 - 1.1838902 | 0.44003 |
| Water-depth | -0.07316 | 0.03113 | -1.341682e-01 - -1.215556e-02 | 0.0187 * |
| Soil type | -0.51384 | 0.31513 | - 1.131483e+00 - 1.038082e-01 | 0.1030 |
| Competitor snails | -23.64088 | 9561.62528 | -1.876408e+04 - 1.871680e+04 | 0.9980 |
| Water velocity | -0.01207 | 0.02130 | -5.381777e-02 - 2.967123e-02 | 0.5708 |

\*Statistical significance at $p < 0.05$

## 5.0 DISCUSSION

This study presents an extensive survey of *Biomphalaria* snails in East Africa with sampling undertaken from 156 sites across Kenya, Uganda and Tanzania. It was established that *Biomphalaria* snails are widely distributed across East Africa and that the snails have a preference for streams (50% of all of the sites where the snails were found were streams), followed by rivers at 20.6%. *Biomphalaria* snails were also found in lake habitats, with *Biomphalaria* snails collected from Lake Albert, Lake Babati and Lake Tanganyika. The snails were also found in irrigation schemes, dams and springs. In Kenya, the snails were found around the Lake Victoria basin (Kisumu County, Busia County, Siaya County, Kisii County, Homa Bay County and Nyamira County); at Mwea irrigation scheme, Machakos County, Kitui County and Makueni County. In Uganda, the snails were found around the Lake Victoria basin, at Lake Albert, Masindi Port, Kyenjojo, Pakwach, Kole district and Soroti district. In Tanzania, the snails were found in Babati district, Kilimanjaro region, Iringa region, Tabora region and Kigoma region. Transmission of schistosomiasis is not possible in areas where there is absence of snail intermediate hosts and therefore, understanding the distribution of the snails has a role to play in prevention and control of the disease. This knowledge on the distribution of *Biomphalaria* snails in East Africa can support surveillance responses in terms of detecting new transmission foci and potential schistosomiasis hotspots.

Ecological and physicochemical parameters have an influence on the distribution of *Biomphalaria* snails in East Africa. Temperature was established to be a crucial factor as far as geographical distribution of the snails is concerned with water temperature established to be the most significant parameter that influences the distribution of the snails (p-value= 0.00184). Temperature as a climatic factor is very important for the survival of *Biomphalaria* snails (Yang *et al*., 2018). Sites where less than 20 snails were found had a temperature range of between 19.1°C to 21.3°C. On the other hand, sites where more than 20 snails were found had a temperature range of between 23.2°C to 26.1°C. This suggests that temperature has a role to play in the abundance of *Biomphalaria* snails in aquatic habitats. According to Yang *et al*. (2018), ideal temperature is important for the fecundity of snail intermediate hosts. Sturrock (1965) states that *Biomphalaria* snails are active at temperatures of between 18°C and 32°C while reproduction and survival of the snails is optimum at a temperature range of between 20°C and 26°C. Optimum temperature is also important as far as the production of cercariae is concerned (Yang *et al*., 2018). Previous studies suggest that *Biomphalaria* snails are not as temperature tolerant as *Bulinus* snails and high temperatures discourage their habitation (Sturrock, 1965). Our findings reveal that sites that had temperatures of above 27°C had low snail densities compared to sites that had temperatures of around 25°C. This explains why *Bulinus* snails have previously been found in arid and semi-arid areas whereas *Biomphalaria* snails are rarely found in such environmental conditions (Sturrock, 1965). Our findings corroborate with a study by Ofulla *et al*. (2013) which avers that *Biomphalaria* snails are less tolerant of temperatures above 27°C. Habib *et al*. (2021) asserts that temperature is an important factor for *Biomphalaria* snails’ reproduction and survival and that temperature changes may alter the breeding, survival and distribution of the snails. A positive correlation between water temperature and *B. sudanica* has also been reported previously (Kazibwe *et al*., 2006). Temperatures higher than optimal values negatively interfere with egg production and development of reproductive organs (Habib *et al*., 2021).

Water-depth was also established to be of statistical significance in relation to *Biomphalaria* snail abundance (p-value=0.0187). Snail abundance decreased with increase in water depth. High snail densities were found in Kakulutuine stream, a shallow stream less than 0.3 m deep and the high snail density could be attributed to the shallowness of the stream. A study by Hamman *et al*. (2000) asserts that water depth is an important factor as far as distribution of snail intermediate hosts is concerned and that there is a correlation between shallowness of a water body and the abundance of the intermediate hosts. In addition, Ofulla *et al*., (2013) showed a negative correlation between snail density and water depth in aquatic habitats of Lake Victoria basin. It is worth pointing out that water is an important factor for the survival of snails but too much of it decimates snail populations (Manyangadze *et al*., 2021).

Presence of vegetation in aquatic habitats has a role to play in the distribution of freshwater snails (Lodge *et al*., 1987). Although we did not find statistical significance between vegetation and snail abundance (p-value=0.21828), previous studies have reported that aquatic plants play an important role in the distribution and habitation of snails because they provide the necessary conditions for feeding, oviposition, breeding as well as shelter (Gallardo *et al*., 2017). The types of vegetation that were found in close proximity to *Biomphalaria* snails included submerged vegetation, floating vegetation, phytoplankton, *Nymphaea caerulea, Nasturtium officinale, Enhydra fluctuans, Colocasia esculenta*, algae and *Maranta arundinacea. Enhydra fluctuans* has been associated with *B. sudanica* snail abundance along the shores of Lake Victoria in Mbita, Kenya (Odero *et al*., 2019). Shallow vegetation has also been associated with the abundance of *B. sudanica* snails in Mwanza region, on the shores of Lake Victoria, in Tanzania (Gouvras *et al*., 2017). According to Kloos *et al*. (2014), presence of *Nasturtium officinale* is directly linked to the abundance of *B. glabrata* snails. On the other hand, *Nymphaea caerulea* has been associated with the abundance of *Biomphalaria* snails (Ofulla *et al*., 2013). A study by Thomas (1987) suggests that *Biomphalaria* snails can utilise submerged macrophytes as a nutritional source.

Soil type has an influence on snail abundance (Ofulla *et al*., 2013). Despite the fact that we did not find statistical significance between soil type and snail abundance (p-value=0.21828), we noticed that muddy soils promoted the habitation of the snails. Muddy soils have been noted to be important as far as survival of *Biomphalaria* snails is concerned because they protect snails from desiccation during hot and dry seasons (Lydig *et al*., 2009). Sandy soil is not conducive for the habitation of *Biomphalaria* snails and low snail abundance was found in water bodies rich in sandy soil. An example is Tulimuimbu stream, in Eastern province, Kenya that had lower snail abundance. Lydig *et al*. (2009) posits that rich black cotton soils that are rich in organic matter are favourable for the habitation of *Biomphalaria* snails. According to Ofulla *et al*. (2013), the substratum in any given habitat has a role to play in the availability of snails and clay soil and silt constitute very important substrata favourable for the survival of *Biomphalaria* snails. Therefore, absence of *Biomphalaria* snails in a given location could be due to the lack of the right substrata.

Water velocity has been established to have an influence on the distribution and abundance of freshwater snails (Rabone *et al*., 2019; Ntonifor & Ajayi, 2007; El-Deeb *et al*., 2017). In spite of the fact that we did not find statistical significance between water velocity and snail abundance (p-value= 0.5708), it has been reported previously that *Biomphalaria* snails are not found in sites with water velocities greater than 30cm/s (Appleton, 1978). We observed that water velocity higher than 30cm/s negatively correlated with *Biomphalaria* snails abundance. Water velocities higher than 30cm/s are a limiting factor as far as snail abundance is concerned. According to Appleton (1978), water velocities have a tendency to sweep *Biomphalaria* snails from one location to another, mostly during the rainy season. Floods are a bottleneck that has been confirmed to have a direct impact on the availability and distribution of *Biomphalaria* snails (Habib *et al*., 2021).

Competitor snails of *Biomphalaria* snails are regarded as biotic factors that affect the distribution of *Biomphalaria* snails (Pointier *et al*., 1993). While we did not find statistical significance between the presence of *Melanoides tuberculata* (a competitor snail) and the abundance of *Biomphalaria* p-value=0.9980), previous studies affirm that *M*. *tuberculata*, a highly invasive snail species is very effective in displacing *Biomphalaria* snails (Geshaw *et al*., 2008 and Pointier, 1993). *Biomphalaria* snails were absent in most of the sites where we found *M*. *tuberculata* in our study except at Ntoroko beach (Lake Albert). Competitor snails of *Biomphalaria* snails in East Africa would therefore seem to have an effect on the distribution of *Biomphalaria* in the region albeit not significant in our glmmTB analyses. Competitor snails have been used in schistosomiasis control by eliminating the snail intermediate host thus interrupting the transmission of *Schistosoma* parasites. Natural enemies of *Biomphalaria* snails such as *Marisa* snail that competes with *Biomphalaria* snails for food and also eats their eggs has been used to control snail populations of *Biomphalaria* snails. *Marisa* species are widely distributed in South America, Africa, Central America and the Caribbean (Muller, 1975). Introduction of *Marisa* snails in new habitats occupied by *Biomphalaria* snails has played a major role in decimating *Biomphalaria* snail populations (Muller, 1975).

## CONCLUSION

This study showed that *Biomphalaria* snails are widely distributed in East Africa and its distribution is influenced by ecological and physicochemical factors. Temperature and water depth were established as the most important factors that influence the geographical distribution of the snails. Other factors that influence the distribution of the snail intermediate hosts in East Africa, albeit not statistically significant include vegetation, algae, pH, water-velocity, soil type and presence of competitor snails. This study provides updated information on the distribution of *Biomphalaria* snails in East Africa and contributes towards the body of knowledge on schistosomiasis and the snail intermediate hosts. Although *Biomphalaria* snail sampling was conducted extensively in East Africa, further sampling should be conducted in future because of the seasonality of the water bodies that are inhabited by the snails and the ravages caused by floods and droughts to those very habitats.

